# Long-read sequencing reveals the complex splicing profile of the psychiatric risk gene *CACNA1C* in human brain

**DOI:** 10.1101/260562

**Authors:** Michael B Clark, Tomasz Wrzesinski, Aintzane B Garcia, Nicola A. L. Hall, Joel E Kleinman, Thomas Hyde, Daniel R Weinberger, Paul J Harrison, Wilfried Haerty, Elizabeth M Tunbridge

## Abstract

RNA splicing is a key mechanism linking genetic variation with psychiatric disorders. Splicing profiles are particularly diverse in brain and difficult to accurately identify and quantify. We developed a new approach to address this challenge, combining long-range PCR and nanopore sequencing with a novel bioinformatics pipeline. We identify the full-length coding transcripts of *CACNA1C* in human brain. *CACNA1C* is a psychiatric risk gene that encodes the voltage-gated calcium channel Ca_V_1.2. We show that *CACNA1C*’s transcript profile is substantially more complex than appreciated, identifying 38 novel exons and 241 novel transcripts. Importantly, many of the novel variants are abundant, and predicted to encode channels with altered function. The splicing profile varies between brain regions, especially in cerebellum. We demonstrate that human transcript diversity (and thereby protein isoform diversity) remains under-characterised, and provide a feasible and cost-effective methodology to address this. A detailed understanding of isoform diversity will be essential for the translation of psychiatric genomic findings into pathophysiological insights and novel psychopharmacological targets.

## Introduction

Genomic studies have identified numerous common single nucleotide polymorphisms (SNPs) that are robustly associated with psychiatric disorders; the challenge is now to understand the underlying pathophysiological mechanisms^1^. Most of these SNPs are non-coding^2–4^, implying that they mediate disease associations by influencing aspects of RNA expression. Of the various mechanisms of RNA regulation, the possibility that psychiatric risk SNPs might influence RNA splicing is particularly appealing, given its exquisite, cell type-specific regulation and its key role in determining neuronal properties^5^. Consistent with this hypothesis, at a global level, cellular studies emphasise RNA splicing as a key mechanism mediating the effect of disease-associated, non-coding variants in complex disorders, including schizophrenia^6^. Similarly, in human brain, genomic regions associated with schizophrenia are enriched for genes that show differential isoform usage across neurodevelopment^7^, implicating many schizophrenia-associated risk loci in the regulation of the expression of specific RNA transcripts. Consistent with these findings, examples of associations between psychiatric risk-associated loci and the abundance of novel, alternatively spliced transcripts are beginning to emerge^8–10^, complementing numerous reports of altered splicing patterns in the brains of psychiatric cases, compared to controls^11–13^.

Splicing varies extensively across human development and aging^7^, and between tissues. The brain exhibits one of the highest levels of splicing diversity and prominent use of tissue specific exons, microexons and splicing factors^14–18^. However, despite extensive efforts to improve the annotation of the human genome^19^, the alternative splicing patterns of many genes remain poorly described, especially in brain, and novel coding exons (many of which are enriched within brain-expressed genes^20, 21^) and transcripts are continually being discovered. This is due to the limited availability of high-quality, post-mortem human tissue and to technical limitations associated with standard approaches: most rely on short-read RNA-Seq and the reconstruction of fragmented sequences making the disambiguation of full-length transcripts difficult, particularly for large and complex genes. Benchmarking studies have shown that short-read RNA-Seq methodologies are unable to accurately reconstruct and quantitate the majority of transcript and protein isoforms^22, 23^.

The relative lack of knowledge about the splicing of individual human genes is typified by *CACNA1C*. *CACNA1C* encodes the Ca_V_1.2 voltage-gated calcium channel (VGCC) alpha_1_ subunit and is a leading genomically-informed candidate gene for multiple psychiatric disorders^2, 3, 24, 25^. However, it is not known how its expression and/or splicing is altered in illness, or in association with genetic risk for it^26^. The *CACNA1C* gene is large (cDNA <13433 nucleotides [nt], 2209 amino acids [aa]) and complex, with at least 50 annotated exons and 31 predicted transcripts (Gencode 27; Figure 1A)^17^. Its size and complexity makes accurate transcript identification and quantification by standard gene expression measures extremely difficult; consequently, the true full-length protein sequence of isoforms remains unclear. Information on the transcript diversity of *human* neuronal VGCC subunits is sparse: most studies have focused on rodents^18^, or human cardiac tissue^27, 28^; the sole study examining VGCC splicing in human brain used a single adult sample^29^. This information is significant, both in terms of understanding pathophysiological mechanisms and for the development of novel pharmacological agents, because *CACNA1C* encodes multiple alternatively spliced transcripts, which result in functionally and pharmacologically distinct channels^30, 31^.

**Figure 1:**
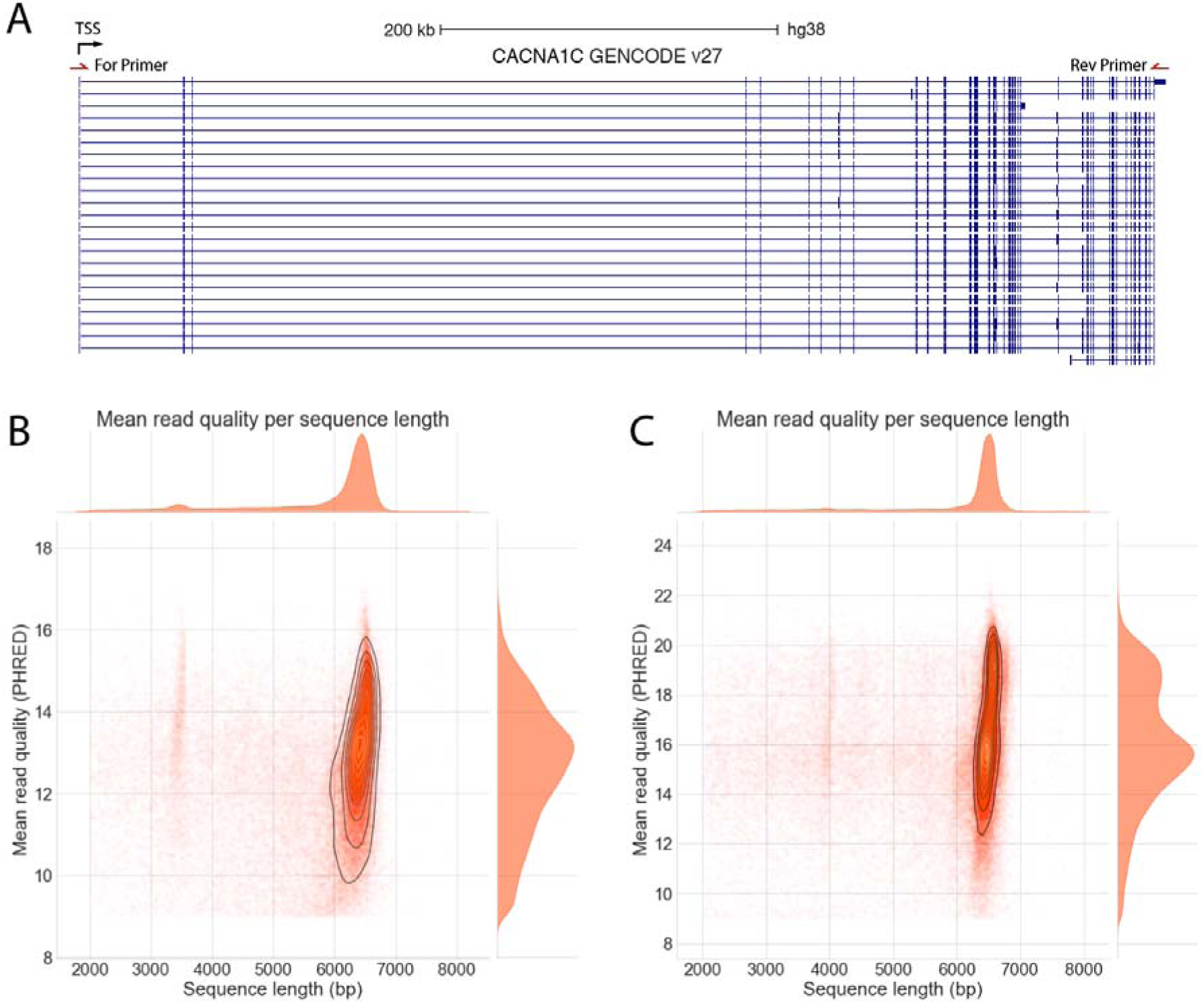
Amplicon sequencing of CACNA1C. A) UCSC genome browser screenshot of CACNA1C isoforms annotated in GENCODE V27. All transcripts in “Basic” annotated set shown. Black arrow shows direction of transcription. TSS: Transcription start site. Position of Forward and Reverse long-range PCR primers in first exon and universal portion of final exon shown. B&C) Length vs Quality of all 2D pass reads from B) Run 1 C) Run 2. Most reads are the full-length CACNA1C CDS. The presence of the 3.5kb positive control CDS spike-in can be seen in Run1. Visualisation limited to reads between 2 and 8 kb, encompassing >98% of pass reads in each run.

To address these unknowns, we developed a straightforward and cost-effective approach using a long-read sequencing technology^32–34^. We combined long-range RT-PCR with long-read nanopore sequencing and a novel bioinformatics pipeline to characterise full-length *CACNA1C* coding sequences (CDSs) in post-mortem human brain. We show that the transcript structure of human *CACNA1C* is substantially more complex than currently appreciated, with numerous novel transcripts and isoforms containing unannotated exons, novel splicing junctions, and in-frame deletions that are predicted to alter the protein sequence and function.

## Methods

### Sample preparation and sequencing

Post-mortem brain tissue from three adult donors in the Lieber Institute for Brain Development repository was obtained from the Maryland Medical Examiner’s Office^35^. Subjects with evidence of neuropathology, drug use (other than alcohol), or psychiatric illness were excluded. Demographic information is presented in Supplementary Table 1.

Full methodological details are presented in the Supplementary Information. Briefly, RNA was extracted from cerebellum, striatum, dorsolateral prefrontal cortex [DLPFC], cingulate cortex, occipital cortex and parietal cortex and reverse transcribed. We selected these brain regions to investigate transcript diversity in cortical and subcortical regions known to be transcriptionally and functionally diverse^36^ and to investigate to what extent the *CACNA1C* transcript profile differs between cortical regions. Full-length CDSs (from the first to last exon [focussing on ‘Exon1B’ the start exon most abundant in brain^30^]) were amplified using PCR, and barcoded and sequenced (across two runs; six samples common to both runs) using Oxford Nanopore Technology’s MinION.

### Data analysis and code availability

Our novel data analysis pipeline is available from: https://github.com/twrzes/TAQLoRe. 2D pass reads (i.e.: those with a q-score of >=9) with an identified barcode were mapped to the transcriptome (hg38, ENSEMBL v82) using a HPC version of BLAT (pblat-cluster v0.3, available from http://icebert.github.io/pblat-cluster/^37^) and to the *CACNA1C* meta-gene containing known exonic sequences using GMAP version 2017-04-24^38^. Reads mapping uniquely to *CACNA1C*, and with at least 50% of their length mapped were retained for further analysis.

To identify potential novel exons, we mined the alignments of the reads to the transcriptome for inserts of at least 9 nucleotides in length located at the junctions between annotated exons. To further characterize candidate novel exons, we mapped the corresponding reads to the genome using LAST v926^39^. We retained candidate exons located within the expected intronic sequences, and at least 6 nt away from existing exons. Novel exonic sequences were subsequently introduced to the *CACNA1C* metagene (concatenation of all exons) and reads were mapped again to this new model using GMAP to enable further characterization of alternative splicing and quantification of transcript expression.

To characterize novel exon junctions, we mapped all the reads to the human genome (hg38) using LAST v926^39^. We identified all exon junctions that showed perfect and contiguous mapping (no sequencing errors nor insertion/deletions) of the read to the genome and used canonical acceptor and donor splice sites. We then used this comprehensive set of exon junctions to correct reads with sequencing errors at mapping breakpoints by selecting the closest canonical splice site to the breakpoint.

To annotate transcripts that include novel exons and/or junctions, we parsed the CIGAR string from the alignment to the metagene, to identify the combination of exon junctions supported by each read. We subsequently clustered splicing patterns to annotate a unique set of isoforms for *CACNA1C* and enable their expression quantification. We applied a similar approach to annotate the transcripts containing novel splice sites. We filtered exons and transcripts of low abundance (i.e. supported only by one read), and clustered transcripts together to call only their longest possible variants.

Transcript expression both on exon and splice site levels was quantified using the number of reads supporting the transcript model. Reads mapping to multiple transcripts were down-weighted according to the number of transcripts they could be mapped to. Read counts were normalized across libraries using the trimmed Mean of M-values normalization method (TMM)^40^. For visualisation, all normalized counts were log_10_-transformed. Expression heatmaps and PCA plots were generated using R statistical language^41^, by heatmap3^42^ and ggplot2 libraries^43^, respectively. To normalise for the difference in sequencing depth between samples, we downsampled all libraries to match that with the smallest sequencing depth and recomputed transcript expression patterns.

### Quality control

A subset of novel exons and novel exon junctions found in *CACNA1C* by nanopore sequencing (Supplementary Table 2) were confirmed by PCR targeting the novel sequence followed by Sanger sequencing (See Supplementary Information for details).

RNA extracted post mortem shows variable degradation, and its quality is usually assessed using the RNA integrity number (RIN). In order to identify the minimum RIN required for reliable amplification of the full-length CDS of *CACNA1C*, RNA samples were artificially degraded (Supplementary Figure 1). *CACNA1C* CDS amplification in these artificially-degraded samples was assessed alongside that from striatal RNA samples of varying RINs from three adult donors (Supplementary Table 3).

## Results

We successfully amplified and sequenced the full length ∼6.5kb *CACNA1C* CDS (encompassing the full intron chain of *CACNA1C*) from all samples (Figure 1A). A minimum RIN of 6 is required, and a RIN of >7 is optimal, for amplification of full-length *CACNA1C* (Supplementary Figure 1). Sequencing run 1 produced 112024 reads, including 52994 (47%) 2D pass reads with an identified barcode. Run 2 produced 126314 reads, including 83221 (64%) 2D pass reads with an identified barcode (Supplementary Tables 4 and 5). Most pass reads were full length, with the updated flowcell used in Run2 providing higher quality reads with a lower error rate (Figure 1). All analyses were performed using the 2D pass barcoded reads, hereafter referred to simply as “reads”.

### Long-range amplicon sequencing reveals many novel CACNA1C exons and isoforms

We developed a bespoke alignment and mapping pipeline to maximise the transcript information obtained from nanopore sequencing reads, including the identification of novel exons, acceptor and donor splice sites, and splice junctions (Supplementary Figure 2). Because of *CACNA1C’s* complexity, we used two complementary approaches to identify transcripts: exon-level and splice-site level analyses. These combined approaches identified a total of 251 unique *CACNA1C* transcript isoforms present in human brain, 241 of which are novel, including the use of novel exons, novel splice sites and junctions. The identity of these transcripts will be made available as a track hub that can be viewed on the UCSC Genome Browser on publication.

#### Exon-level analysis

We annotated a total of 39 potential novel exons within the *CACNA1C* locus, of which 38 were identified in at least 2 individuals or tissues and supported by at least 5 nanopore reads in each library (Figure 2A; Supplementary Data Table 1).

**Figure 2.**
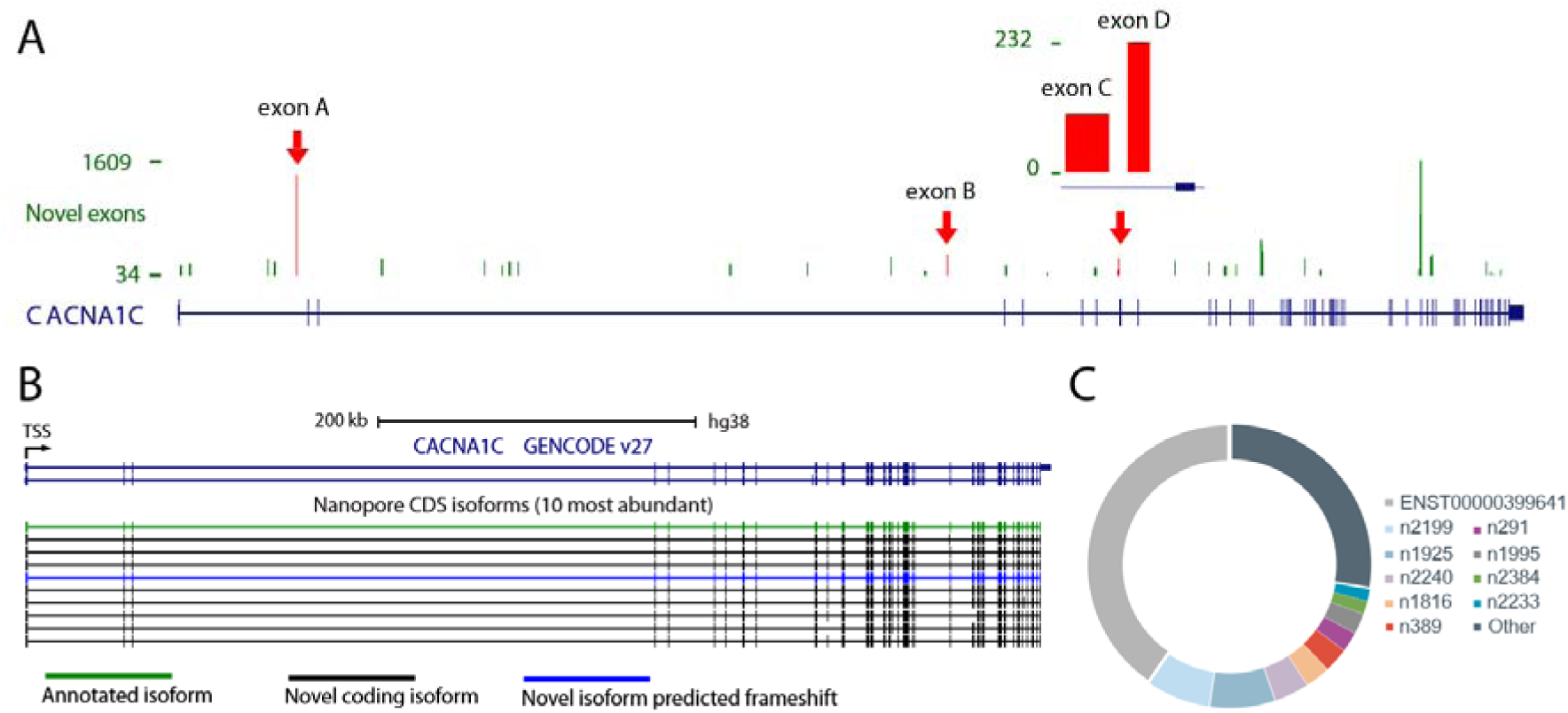
(**A**) Annotation and read count for novel exonic sequences within CACNA1C. Red arrows indicate exons that have been validated (Supplementary Table 2). (**B&C**) Top 10 most abundant CACNA1C isoforms identified in brain using exon-level approach. (**B**) UCSC genome browser screenshot of top isoforms. Colours denote transcript type. (**C**) Proportion of high confidence transcripts reads from the ten most abundant transcripts.

We validated four out of four of the novel exons (selected across the range of abundance) by PCR and Sanger sequencing by confirming the splice junctions between the novel exon and its surrounding annotated exons. We also discovered a fifth novel exon that was spliced in between the targeted novel exon and the nearest known exon. This successful validation of a selection of novel exons provides high confidence that the novel exons identified by nanopore sequencing are real and actively incorporated into *CACNA1C* transcripts.

Integration of these 38 novel exons and the mapping of the reads to the metagene enabled the identification of novel and known (i.e. annotated) transcripts. Novel transcripts incorporate novel exons, and/or novel junctions between annotated exons and/or new combinations of known junctions. To limit the impact of sequencing errors on transcript annotations, we filtered the identified transcripts retaining only those found in at least 2 libraries with a minimum of 24 reads in total (see Supplementary Data Files 2 and 3 for transcript identity and abundance, respectively). We also created a high confidence set of transcripts with an increased minimum threshold of 100 reads (see Supplementary Data Files 4 and 5 for transcript identity and abundance, respectively). Unless otherwise stated all analyses were performed on the high confidence transcripts set. We identified 90 high confidence *CACNA1C* transcripts across the 6 brain regions, including 7 previously annotated (GENCODE v27) and 83 novel (Supplementary Figure 3). Seven of the novel high confidence transcripts contained novel exons (1.8% of high confidence reads in total), while the remaining 76 included previously undescribed junctions and junction combinations.

As expected the most highly expressed transcript was previously annotated (`ENST00000399641`) and is supported by 40.2% (23.7%-50%) of reads on average. In comparison, the most highly expressed novel transcript (*CACNA1C n2199*) represents on average 7.3% (2.7%-25.3%) of all reads (Figure 2B,C). Nine of the top ten expressed transcripts were novel, of which eight are predicted to maintain the *CACNA1C* reading frame, suggesting a number of these most abundant novel transcripts encode functionally distinct protein isoforms (Figure 2B,C). Without any thresholding, 75.5% of all the reads supported previously unannotated transcripts. These results suggest that novel *CACNA1C* transcripts are abundantly expressed as well as highly numerous, and that current annotations are missing many of the most abundant *CACNA1C* transcripts. Without filtering, we found evidence for the expression of only 18 of the current set of 31 annotated transcripts (GENCODE v27).

#### Splice-site level analysis

The exon-level transcript identification approach described above provides a robust and conservative means to identify novel exons and to characterise full-length transcript structure. However, it is relatively insensitive to small-scale variation (of the order of a few amino acids). Therefore, we also implemented a splice junction-level analysis to annotate novel splice sites and junctions in the *CACNA1C* gene model. We used a conservative approach, which relies on the identification of junctions supported by error-free mapping at the junction, and the presence of canonical splice sites. We thereby identified 497 novel splice sites, of which 393 were supported by at least 10 reads. Compared to previously annotated splice sites, novel donor and acceptor splice sites are less used (Supplementary Figure 4). Including these splice sites, and after filtering for transcripts supported by at least 24 reads, we identified 195 transcripts, of which 111 are predicted to be coding (see Supplementary Data Files 6 and 7 for transcript identity and abundance, respectively). 28 of these were identical to transcripts identified by the exon-level analysis. Strikingly, most of the remainder (130) resulted from small-scale (≤ 15 nucleotides) differences to our high-confidence exon-level transcripts, demonstrating the complementarity of these approaches. We validated the presence of specific small-scale deletions (3 and 4 amino acid microdeletions, described below) using Sanger sequencing.

### The expression profile of CACNA1C isoforms differs between brain regions

We examined how *CACNA1C* isoform expression varies across brain regions and between individuals, focussing on transcripts identified using the exon-level approach (similar results were obtained using the splice-site level transcripts [Supplementary Figure 4). We downsampled all libraries to 2,729 (the smallest sequencing depth) prior to normalization. Differences between tissues is the main driver of the observed variation in expression between transcripts. Cerebellum and striatum were distinct from the four regions of the cortex (Figure 3A,B), but expression was similar across individuals, consistent with observations at the transcriptome level^44^. The use of the more permissive filtered set of transcripts for expression estimation (see Methods) further improved the separation between regions, and highlighted potential differences in expression between cortical regions (Supplementary Figures 5 and 6).

**Figure 3.**
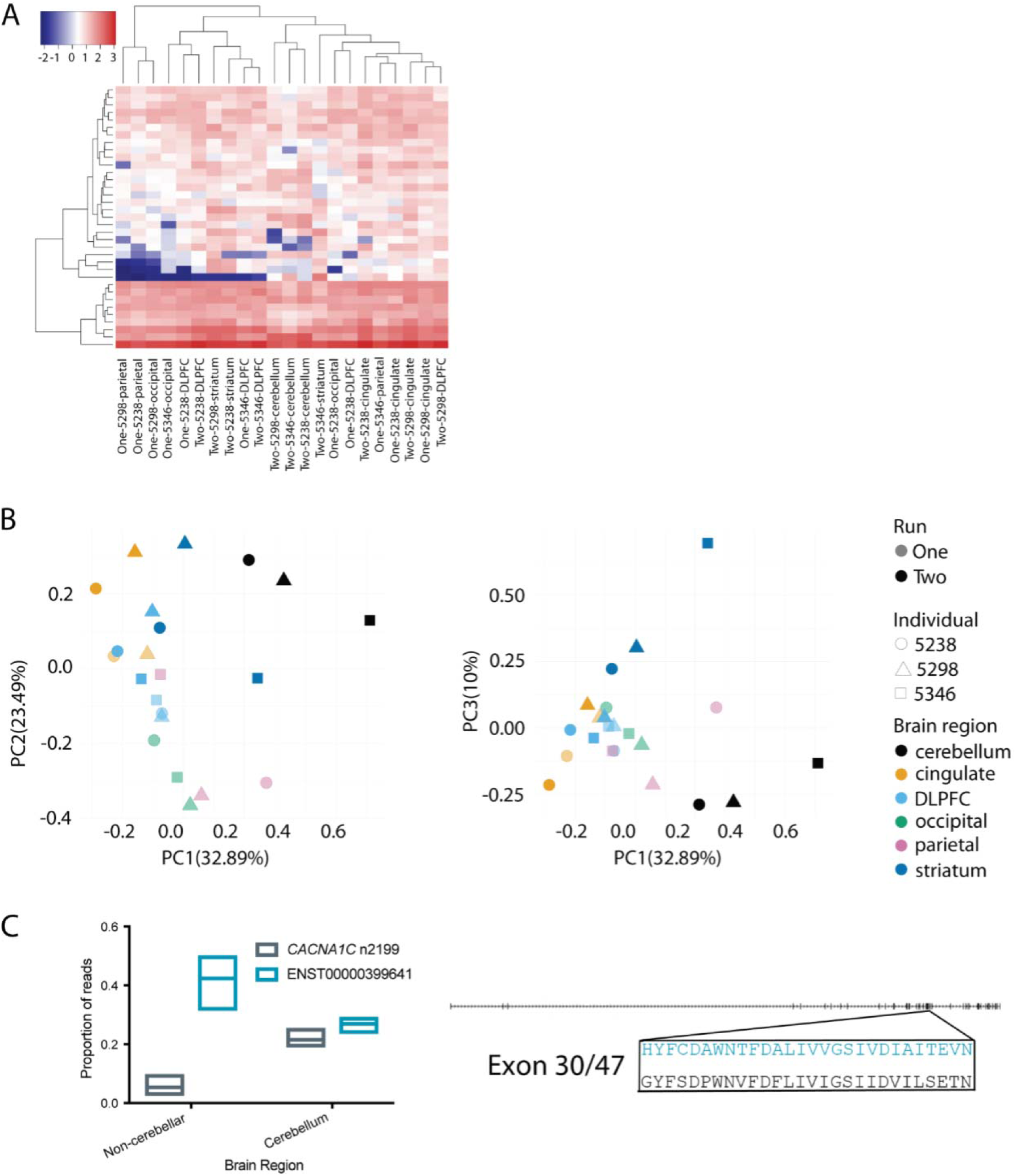
Comparison of CACNA1C isoform expression between individuals and tissues (**A**) Transcript expression levels (TPM) across tissues and individuals. “One” and “Two” denote sequencing runs. (**B**) Principal Component Analysis based on normalised transcript expression. (**C**) Isoform switching of ENST00000399641 and CACNA1C n2199 in cerebellum. Left panel: box plots show minimum to maximum values with line at mean value. Right panel: the sequences of ENST00000399641 (blue) and CACNA1C n2199 (black) differ in the sequences of their 30^th^ exons. The detail shows the amino acid sequences of this exon in the two transcripts.

Although we did not find any tissue-specific transcripts amongst our set of high confidence transcripts, we observed a pronounced transcript expression switch in cerebellum. Outside of the cerebellum *ENST00000399641* was the dominant transcript, whilst in cerebellum, *ENST00000399641* and *CACNA1C n2199* were expressed at similar levels (Figure 3C).

### Predicted impact of novel isoforms on the Ca_V_1.2 protein model

*CACNA1C* encodes the pore-forming Ca_V_1.2 alpha_1_ VGCC subunit. The calcium pore conmprises 24 transmembrane repeats, clustered into 4 domains linked by intracellular loops (Figure 4A). Among the 83 novel exon-level transcripts we identified, 51 potentially encode functional Ca_V_1.2 channels, whilst the remainder are likely noncoding as they contain deletions in critical membrane-spanning regions and/or frameshifts. Notably, putatively coding isoforms represent 87.8% of total high confidence reads, demonstrating that while noncoding transcripts are numerous, it is putatively coding transcripts that represent the vast majority of reads (Supplementary Figure 7).

**Figure 4.**
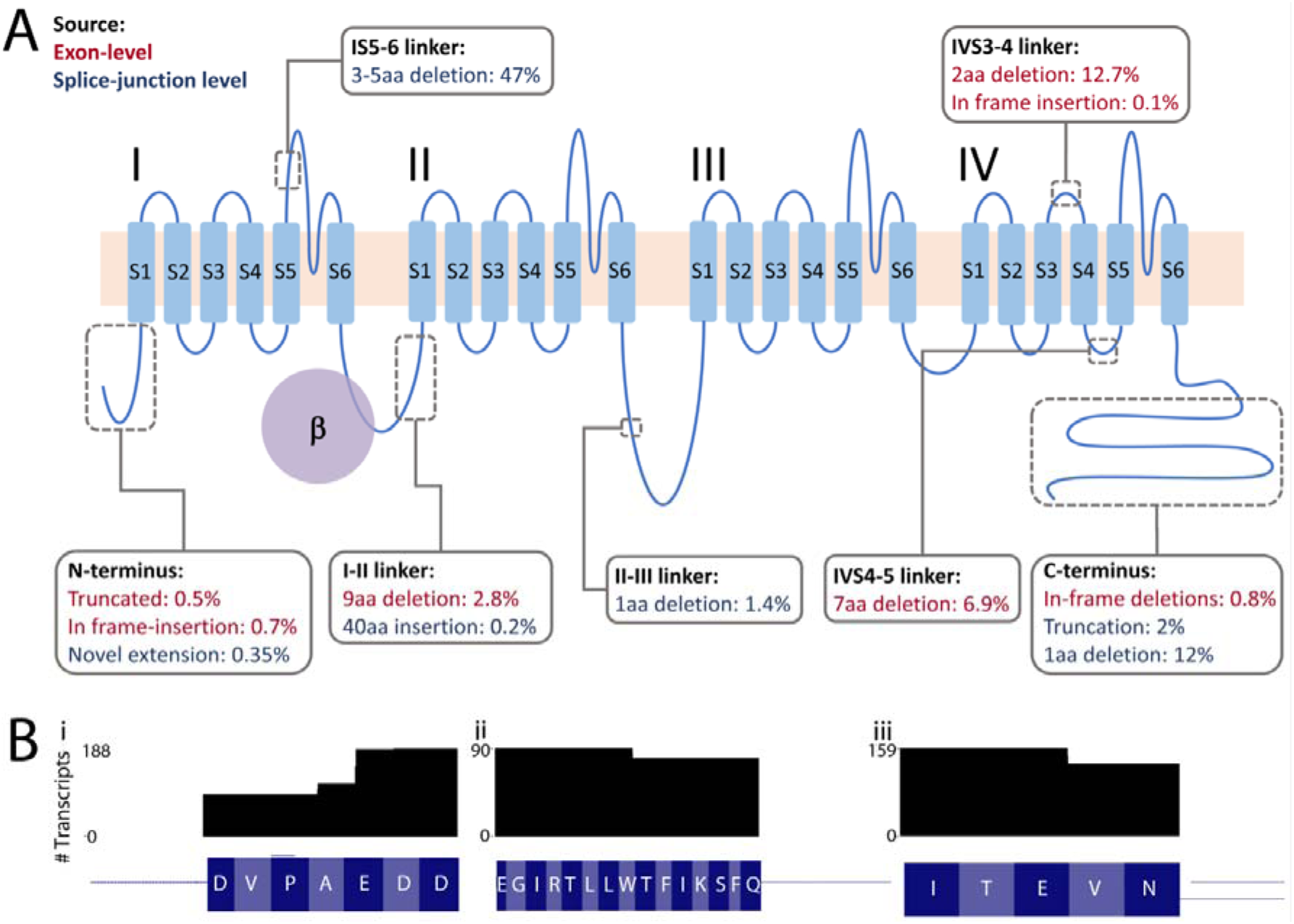
Impact of novel splicing events on the CACNA1C protein model. (A) CACNA1C encodes the primary pore-forming subunit of the Ca_V_1.2. Ca_V_1.2 is formed of 4 domains (I-IV), each comprised of 6 transmembrane domains (S1-S6), which are linked by intracellular loops. The obligate beta subunit binds to the I-II intracellular loop, as shown. Grey boxes indicate the location of novel, in-frame insertions and deletions, discussed in the main text. Values indicate the mean proportion of reads containing each variant. Where variants were identified using both analysis approaches, exon-level counts were used to derive abundance (red text); variants identified only using the splice-site level approach are indicated with blue text. (B) Number of protein isoforms containing 3 microdeletions: (i) in the I-II linker, (ii) in the IV4-5 linker and (iii) the previously-reported microdeletion in the IV3-4 linker.

Around half of putatively coding exon-level transcripts (25 of 51; 26.8% of total coding reads) consist of novel combinations of already annotated exons. The remainder include novel exons and/or deletions (Figure 3B). Many of the novel splicing events are seen across multiple transcripts. For example, five transcripts (2.8% on average, 5.2% maximally, of total coding reads) predict Ca_V_1.2 protein isoforms with an in-frame deletion in the intracellular I-II linker, and seven (6.9% on average, 11.9% maximally, of total coding reads) contain an in-frame deletion in the IVS4-IVS5 linker (Figure 4A). Ten protein isoforms (12.7%, on average, 22.3%, maximally, of total coding reads) - including *CACNA1C n1925*, one of the three most abundant transcripts - include a previously-described microdeletion in the extracellular IVS3-IVS4 linker (Figure 4B)^33^. One also contains a 3aa in-frame insertion in this same linker region, highlighting this region as a ‘hotspot’ for alternative splicing, of interest given the relevance of this linker for determining channel properties^33^.

Several protein isoforms predict alternative N- and C-termini (Figure 4A), which are involved in coupling Ca_V_1.2 signalling to intracellular signalling cascades and calcium-dependent inactivation^31, 45^. Variation in these regions is predicted to affect a small fraction of the Ca_V_1.2 pool, but such protein isoforms may still be of biological significance if they show altered inactivation properties and/or coupling to second messenger systems (Figure 4A).

The splice-site level analysis identified additional sites of variation (Figure 4). Most strikingly, 47% of transcripts included an in-frame microdeletion of 3, 4 or 5 amino acids in the IS5-6 linker (corresponding to amino acids D306-E310 in Uniprot entry Q13936). The 4 amino acid microdeletion was previously described in heart but not brain^29^, but the 3 and 5 amino acid microdeletions are novel (Figure 4B). Their functional impact is unknown but the S5-S6 linker is part of the pore-forming region and these microdeletions fall within a region previously implicated in determining channel conductance^46^.

## Discussion

We combined long-range RT-PCR with nanopore sequencing in order to characterize the full coding sequence from transcripts of *CACNA1C*. To our knowledge, we are the first to have used this approach successfully in human postmortem tissue. The vast majority of *CACNA1C* transcripts are novel, and many of these are abundant. We demonstrate marked differences in *CACNA1C* transcript profiles between brain regions, with the cerebellum in particular showing a notable switch in isoform abundance compared with cortex. Our results demonstrate that *CACNA1C* transcripts are much more diverse than previously appreciated, and emphasise the importance of studying full-length isoforms and of access to high-quality human brain tissue.

Despite annotations of very high quality and active curation^19, 47^, an increasing number of studies report novel protein coding sequences and exons in the human genome. The rapid development of long read sequencing technologies opens the unique opportunity to gain an accurate representation of transcript diversity, as each read encompasses a full transcript. This knowledge is particularly critical for genes with complex models, for example a recent study of DSCAM, identified 18,496 transcripts out of over 19,000 of those theoretically possible^48^. We anticipate that long-read sequencing applications will further help decipher transcript expression and splicing, especially for genes expressed in the brain, which exhibits prominent use of alternative splicing and tissue specific exons^14–18, 49^.

Our study highlights the power of long read sequencing for the annotation and characterisation of alternatively spliced transcripts. We demonstrate a five-fold increase in the number of annotated transcripts for *CACNA1C*, identifying novel exons and deletions within the coding sequence, as well as novel combinations of previously annotated exons. Because of the high quality of the genomic assembly at the *CACNA1C* locus^50^, the novel exons, and splice sites we describe are unlikely to result from mapping errors. Instead, our findings support those from transcriptome analyses that indicate that a significant proportion of gene isoforms in human brain remain unannotated^51^. Supporting their potential importance, a number of the novel transcripts are highly abundant individually, and collectively they encompass the majority of reads. Our finding that the vast majority of reads from novel transcripts maintain the *CACNA1C* reading frame also supports the hypothesis that these are functionally relevant and not simply products of “noisy” splicing. Furthermore, we identified several abundant in-frame deletions that are present in a number of transcripts; if translated these could dramatically impact on Ca_V_1.2’s function in the cell^29, 30, 52^. Notably, a number of our novel predicted protein isoforms include alterations in domains known to be important for determining channel properties and coupling to second messenger systems^29, 31, 53^, thereby providing testable hypotheses as to their predicted functional impact. Now that the transcript structure of *CACNA1C* is clearer, it will also be of interest to examine the *cumulative* effect of functional variation across the Ca_V_1.2 protein on channel function. Conversely, we identified a relatively low number of annotated transcripts in our dataset. This is perhaps unsurprising, given that many of the currently annotated transcripts are likely predictions from ESTs and incomplete cDNAs. Taken together, these observations demonstrate the importance of long read sequencing for the accurate characterisation of transcript structure and alternative splicing.

The need to understand the diversity of Ca_V_1.2 isoforms may be of clinical and therapeutic relevance. Firstly, characterization of the complement of full-length *CACNA1C* isoforms is a necessary first step towards understanding how its transcript profile might be altered by a risk variant or disease state. Secondly, calcium channel blockers, which have Ca_V_1.2 as one of their primary targets, are licensed for cardiovascular indications and have possible utility as a therapeutic strategy for psychiatric disorders^54–57^. Since the Ca_V_1.2 proteins that arise from *CACNA1C* splicing show differential sensitivity to the existing calcium channel blockers^52^ it may be possible to selectively target disease-relevant *CACNA1C* isoforms and/or those that are differentially expressed in the brain vs. the periphery, to provide novel psychotropic agents that are both more potent and are freer from peripheral side effects.

Our results indicate differences in the abundance, but not the identity, of *CACNA1C* transcripts between brain regions. Utilising our more conservative exon-level analysis, in most regions there is a single major *CACNA1C* isoform, with levels of expression almost five-fold higher than the second most highly expressed transcript. However, in the cerebellum there is a switch in transcript abundance, such that the two most abundant transcripts are expressed at similar levels to one another. The consistent nature of this switch in different individuals supports the hypothesis that this is a regulated switch in expression. In contrast, our analysis demonstrated only minor differences in the transcript profile between individuals (Figure 3B). Given these findings, we are therefore confident that our analysis is likely to have identified most common and abundant *CACNA1C* transcripts present in adult human brain. However, we anticipate that the relative abundance of these transcripts will likely differ between individuals. Our splice site-level anlaysis found broadly the same results, although there was no longer a single dominant isoform in each region, largely due to variation in the IS5-6 linker. While we have validated these variations, increased certainty in their abundance and impact on overall transcript structure will likely require further improvements in exon boundary identification with nanopore sequencing. The details of the *CACNA1C* transcript structure revealed here will facilitate the mining of existing large-scale RNA-Seq datasets to investigate these differences between individuals (and between subgroups based on e.g. genotype or disease status).

In summary, our findings demonstrate the utility of long-range amplicon sequencing for the identification and characterisation of gene isoform profiles. More specifically, they demonstrate that the human brain *CACNA1C* transcriptional profile is substantially more complex than currently appreciated. Our approach focused on annotated 5’ and 3’ ends; thus, there is a need for further in depth investigation, e.g. using RACE and capture sequencing. Understanding the functional consequences of this isoform diversity will advance our understanding of the role of VGCCs in the human brain and their involvement in psychiatric disorders^2–4^. Finally, some of the novel isoforms we present here may prove novel and selective therapeutic targets for these disorders^54^.

## Acknowledgements

We are grateful to Li Chen and Arne Mould for technical assistance. This research was supported by a Wellcome Trust [201879/Z/16/Z] award to MBC and a UK Medical Research Council [MR/P026028/1] award to EMT. The authors wish to acknowledge the following funding sources: MBC is supported by an Australian National Health and Medical Research Council (NHMRC) Early Career Fellowship [APP1072662]. EMT is supported by a Royal Society University Research Fellowship. WH and TW are supported by the BBSRC, Institute Strategic Programme Grant [BB/J004669/1], BBSRC Core Strategic Programme Grant [BB/P016774/1]. This study was supported by the National Institute for Health Research Oxford Health Biomedical Research Centre. The human brain tissue repository is supported by the Lieber Institute for Brain Development. The views expressed are those of the authors and not necessarily those of the NHS, the NIHR or the Department of Health.

The contents of the published material are solely the responsibility of the administering institution, a participating institution, or individual authors and do not reflect the views of NHMRC.

## Author contributions

MBC and EMT conceived the study. MBC, ABG, PJH, NAH, TW, WH and EMT designed experiments. MBC, ABG and EMT performed experiments and nanopore sequencing. ABG and NAH performed PCR validations. JEK, TH and DRW provided materials. TW and WH wrote the analysis pipeline and performed informatic analyses. MBC, NAH, ABG, TW, WH and EMT wrote the manuscript.

## Data Availability

Nanopore sequencing reads generated and analysed in the present study are available from ENA (*Accession pending*). Transcript isoforms will be made available as supplementary data and a UCSC Genome Browser Track Hub on publication.

Supplementary information is available at MP’s website

## Supplementary Information

### Supplementary Methods

All molecular biology protocols were conducted according to manufacturer’s recommendations unless specifically noted otherwise.

#### RNA extraction and RT-PCR

RNA was extracted from cerebellum, striatum, dorsolateral prefrontal cortex [DLPFC], cingulate cortex, occipital cortex and parietal cortex. Tissue was disrupted with the TissueLyser LT (Qiagen, UK) using Rnase-free 5mm stainless steel beads in QIAzol Lysis Reagent (Qiagen). RNA was extracted using the Qiagen RNeasy Lipid Tissue Mini Kit (Qiagen, UK) and eluted in RNase-free water. RNA concentration and RNA integrity (RNA Integrity Number equivalent; RINe) were measured using a Tapestation 2200 (Agilent, UK). All samples had a RINe of >7.0. RNA (1µg per sample) was converted into cDNA with GoScript™ Reverse Transcriptase (Promega, UK) using oligoDT priming.

Full-length CDS amplicons of *CACNA1C* were obtained by PCR using primers located 5’ to the translation start site in Exon 1B and 3’ to the stop codon in the 3’ untranslated region. Specifically, *CACNA1C* was amplified from cDNA equating to 125ng RNA template from the eighteen human brain samples using PrimeSTAR GXL DNA Polymerase (Takara). Tailed primers were designed to amplify the ∼6.5kb *CACNA1C* coding sequence and included sequence to allow further amplification and barcoding. Forward and Reverse primers were tttctgttggtgctgatattgcCATTTCTTCCTCTTCGTGGCTGC and acttgcctgtcgctctatcttcCCAGGTCACGAGAACAGTGAGG, respectively (*CACNA1C* sequence in capital letters). Amplification was conducted for 25 cycles of 98°C for 10 sec; 57°C for 15 sec; 68°C for 7 min. PCR products were separated on a 1.5% agarose gel (visualised with GelGreen, Biotium, UK). Full-length CDS (∼6.5kb) products were excised and purified using the Qiagen Gel Purification Kit (Qiagen, UK). DNA was barcoded using the Oxford Nanopore Technologies (ONT) PCR Barcoding Kit (EXP-PBC001) using PrimeSTAR GXL DNA Polymerase and 12 cycles of PCR amplification, as described above. PCR products were purified with the Qiagen PCR Purification Kit (Qiagen, UK) and quantified by Qubit (ThermoFisher, UK).

#### Nanopore Sequencing of the full-length CACNA1C CDS

The PCR Barcoding Kit (EXP-PBC001, ONT) contains unique twelve barcodes, hence the 18 samples were split across two flowcells with 12 samples run on each. Six samples were sequenced on both flowcells to allow comparison and normalisation between sequencing runs. Samples and barcodes used in each sequencing run are described in Supplementary Table 4.

For each sequencing run, 132ng of each of the twelve samples was pooled and re-purified using 0.4x Agencourt Ampure XP beads (Beckman Coulter, UK) to concentrate the sample and remove any contaminating small DNA products. The presence of full-length DNA product in the pool was confirmed by 2200 or 4200 TapeStation (Agilent) analysis using gDNA screentape (Supplementary Figure 8) and the sample measured again by Qubit to ensure the presence of sufficient product for nanopore library preparation.

Sequencing libraries were prepared with the 2D Nanopore Sequencing Kit (SQK-LSK208, ONT) using 1µg of *CACNA1C* DNA. DNA end-repair and dA-tailing was performed using the NEBNext Ultra II End-Repair/dA-tailing Module (NEB, UK) in a total reaction volume of 60µl (50µl DNA, nuclease-free water (NFW) and DNA calibration strand (Run1 only), 7 μl Ultra II l Ultra II End-Prep enzyme) for 5 min at 20°C and 5 min at 65°C. Samples were purified using 0.6x Ampure XP beads and eluted in 31µl of NFW. Sample concentrations were measured by Qubit and confirmed retention of >85% of the initial samples. The total volume of *CACNA1C* DNA was utilised for subsequent ligation of sequencing adaptors. Ligation reactions contained 50µl Blunt / TA Ligase Master Mix (NEB), 10μl Adapter Mix 2D, 2 μl Hairpin Adaptor (HPA), 30μl of DNA and 8μl of NFW and incubations were performed at room temperature. After 10 min incubation 1 μl Hairpin Tether (HPT) was added, the sample mixed and incubated for a further 10 minutes. Adapted DNA was purified with MyOne C1 beads (ThermoFisher). 50μl of beads were washed 2x in 100μl bead binding buffer (BBB) and resuspended in 100μl BBB before use. Beads were mixed 1:1 with adapted DNA, incubated on a rotor for 5 min at room temperature to bind DNA to the beads and collected using a magnetic stand. Bead-bound DNA was washed 2x with 150μl of BBB on the magnetic stand and left briefly to dry the beads. Beads were resuspended in 15μl of elution buffer (ELB) and incubated at 37°C for 10 min to elute DNA from the beads. After pelleting the beads on the magnetic stand, 14μl of supernatant containing the adapted DNA library in ELB was collected. Library concentrations were measured by Qubit, recovering >25% of the original starting amount.

Flowcells were primed with a mix of 480 μl Running Buffer with Fuel Mix (RBF) and 520 μl NFW as per manufacturer’s instructions. A loading mix of 35 μl RBF, 25.5 μl library loading beads (LLB), 12 μl *CACNA1C* library and 2.5 μl NFW was prepared, mixed by pipetting and immediately loaded onto the primed “spot-on” flowcell. Sequencing was performed on fresh flowcells; Run 1 utilised a R9.0 (MIN105) flowcell, while Run 2 utilised a R9.4 (MIN106) flowcell. Pore occupancy was >80% indicating a good library and flowcell. Sequencing was allowed to continue until there was a high probability of >1000 high-quality (2D pass) reads from each barcoded sample. Run 1 was base-called with the Epi2Me cloud based service (2D Basecalling plus Barcoding for FLO-MIN105 250bps - v1.125). Run 2 was base-called with Albacore V1.1.0 (FLO-MIN106, SQK-LSK208, barcoding) after the withdrawal of the Epi2Me base-calling service. The absence of a base-calling model for R9.0 in the Albacore software prevents the base-calling of Run1 with this software. Sequencing library metrics (Supplementary Table 5) were generated by Poretools <u>(Loman and Quinlan 2014)</u> and PycoQC <u>(Leger and Leonardi 2018)</u>.

#### Validation of novel exons and junctions

In the case of novel exons, two sets of nested PCRs were conducted: one spanning the upstream exon and the novel exon sequence (‘5’ confirmation’ in Supplementary Table 2), and a second spanning the novel exon sequence and the downstream exon (‘3’ confirmation’ in Supplementary Table 2). Novel junctions were confirmed using a single round of PCR using one primer spanning the novel junction, with the second primer located in the neighboring exon. PCR reactions were performed using iIlustra PuReTaq Ready-To-Go PCR beads (GE Healthcare Life Sciences, UK). PCR reactions were cycled as follows: 95°C for 5 min; 35 cycles of 95°C for 35 sec, 30 sec at annealing temperature (see Supplementary Table 2); 72°C for 2 min, followed by a final 5 min extension at 72°C. Primers sequences and conditions are shown in Supplementary Table 2. PCR products were separated on agarose gels. Products of the predicted size were excised, cleaned using the Qiagen Gel Extraction Kit and ligated into pGEM-T Easy vector (Promega) for Sanger sequencing.

#### Impact of RNA quality on amplification of full-length CACNA1C CDSs

Eleven brain RNA samples (see supplementary table 3) with an average RIN of 8 were pooled and aliquoted into eight samples of 7 μl (875ng each at 125 ng/ μl). Each aliquot was heated at 72°C for different periods of time (0, 2, 5, 10, 20, 35, 60 or 90 minutes) to degrade the RNA. The RINe for each aliquot was then assayed using the Agilent 4200 TapeStation system.

Reverse transcription used 500 ng of RNA and was performed using GoScript™ Reverse Transcriptase. *CACNA1C* amplification was performed as described above, using 5μl of cDNA and 30 cycles of amplification. Forward and Reverse primers were: CATTTCTTCCTCTTCGTGGCTGC and CCAGGTCACGAGAACAGTGAGG, respectively, i.e. identical to those used in the main experiment, but lacking the barcoding tails. Successful amplification of PCR product was confirmed and visualised using both agarose gel electrophoresis and 4200 Tapestation analysis using equal proportions of each PCR. Samples from three additional adult donors were utilised specifically for this experiment to provide the required range of high quality and partially degraded samples.

---

**Supplementary Table 1:** Demographics of tissue donors

| Case ID | Age (years) | Sex | PMI | Ethnicity | Brain RIN | Cause of death |
| --- | --- | --- | --- | --- | --- | --- |
| 5238 | 37.2 | M | 10.5 | AA | 8.9 | Multiple blunt force injuries |
| 5298 | 50.3 | M | 12.5 | AA | 8.7 | Multiple injuries |
| 5346 | 25.1 | F | 24.5 | AS | 8.3 | Multiple injuries |

PMI: postmortem interval; RIN: RNA Integrity Number; M: male; F: female; AA: African American; AS: Asian

**Supplementary Table 2:**
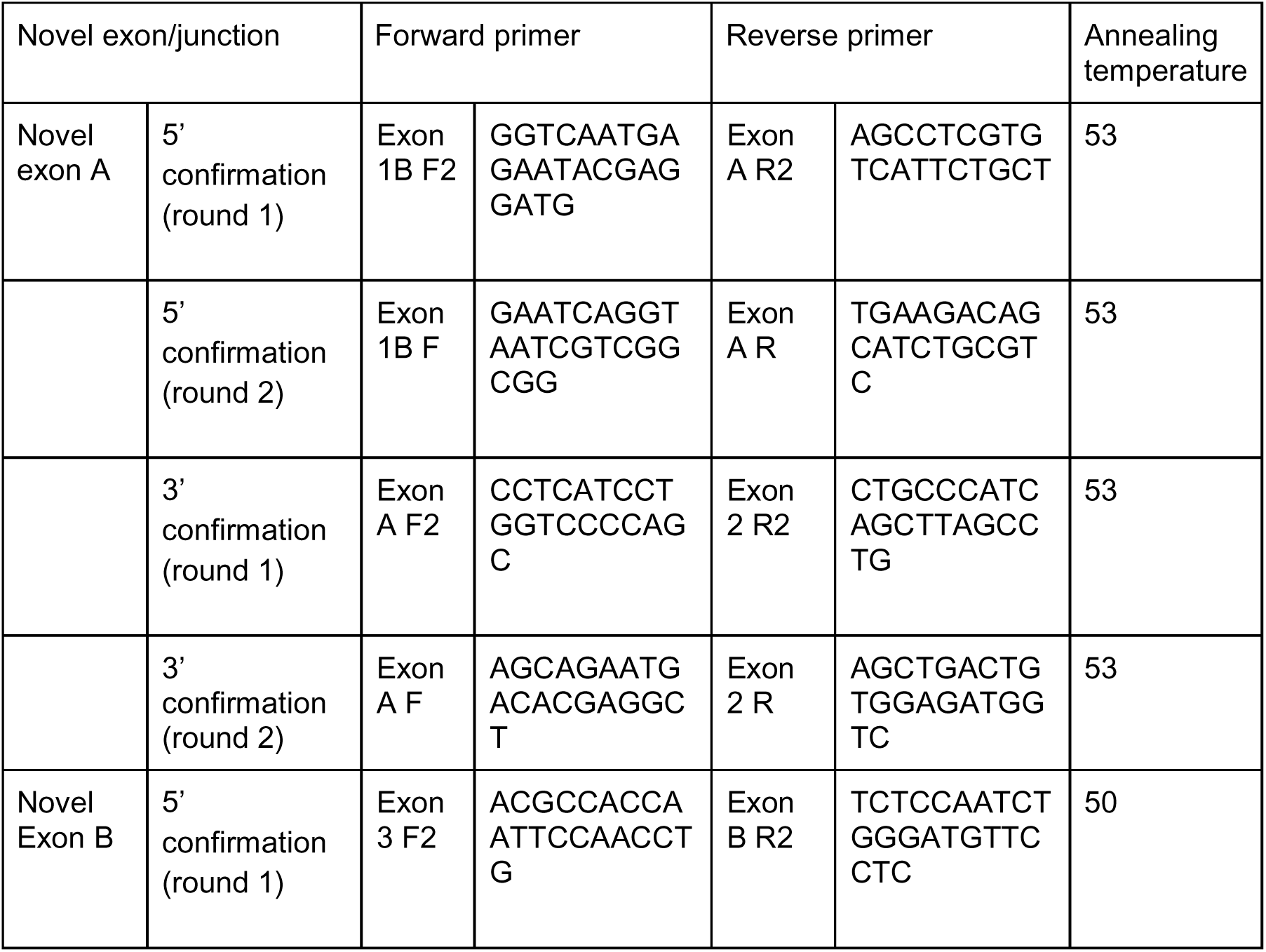
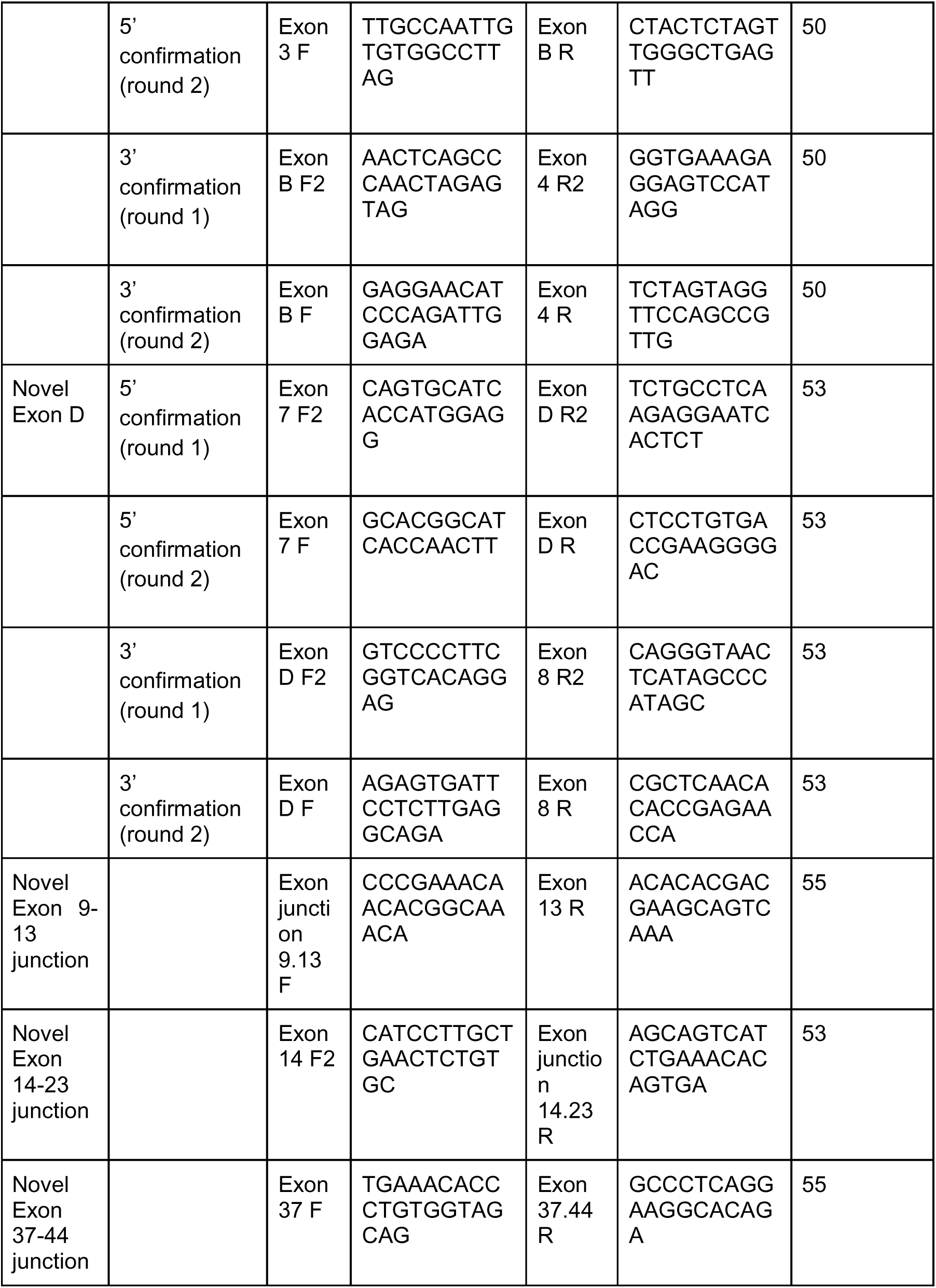
primers and PCR conditions used to confirm novel exons and junctions.

**Supplementary Table 3:** Demographics of additional tissue donors used for RNA degradation study

| Case ID | Age (years) | Sex | PMI | Ethnicity | Brain RIN | Cause of death |
| --- | --- | --- | --- | --- | --- | --- |
| 5244 | 21.2 | M | 28 | AA | 8.7 | Multiple injuries |
| 5579 | 41.9 | F | 13 | CAUC | 6.3 | Multiple injuries |
| 5717 | 51.9 | M | 35.5 | CAUC | 8.9 | Sarcoidosis |

PMI: postmortem interval; RIN: RNA Integrity Number; M: male; F: female; AA: African American; CAUC: Caucasian

**Supplementary table 4:** Samples and pass reads in each sequencing run

| Sample | Barcode | Run | Run1 reads | Run2 reads |
| --- | --- | --- | --- | --- |
| 5238 cingulate | BC001 | 1, 2 | 4836 | 6752 |
| 5238 DLPFC | BC002 | 1, 2 | 7227 | 10484 |
| 5238 occipital | BC003 | 1 | 4630 |  |
| 5238 parietal | BC004 | 1 | 6024 |  |
| 5238 cerebellum | BC003 | 2 |  | 3735 |
| 5238 striatum | BC004 | 2 |  | 8778 |
| 5298 cingulate | BC005 | 1, 2 | 6287 | 9329 |
| 5298 DLPFC | BC006 | 1, 2 | 5660 | 8849 |
| 5298 occipital | BC007 | 1 | 2729 |  |
| 5298 parietal | BC008 | 1 | 2729 |  |
| 5298 cerebellum | BC007 | 2 |  | 6725 |
| 5298 striatum | BC008 | 2 |  | 8116 |
| 5246 cingulate | BC009 | 1, 2 | 1760 | 2421 |
| 5346 DLPFC | BC010 | 1, 2 | 2572 | 3403 |
| 5346 occipital | BC011 | 1 | 4859 |  |
| 5346 parietal | BC012 | 1 | 3681 |  |
| 5346 cerebellum | BC011 | 2 |  | 7609 |
| 5346 striatum | BC012 | 2 |  | 4130 |

**Supplementary table 5:**
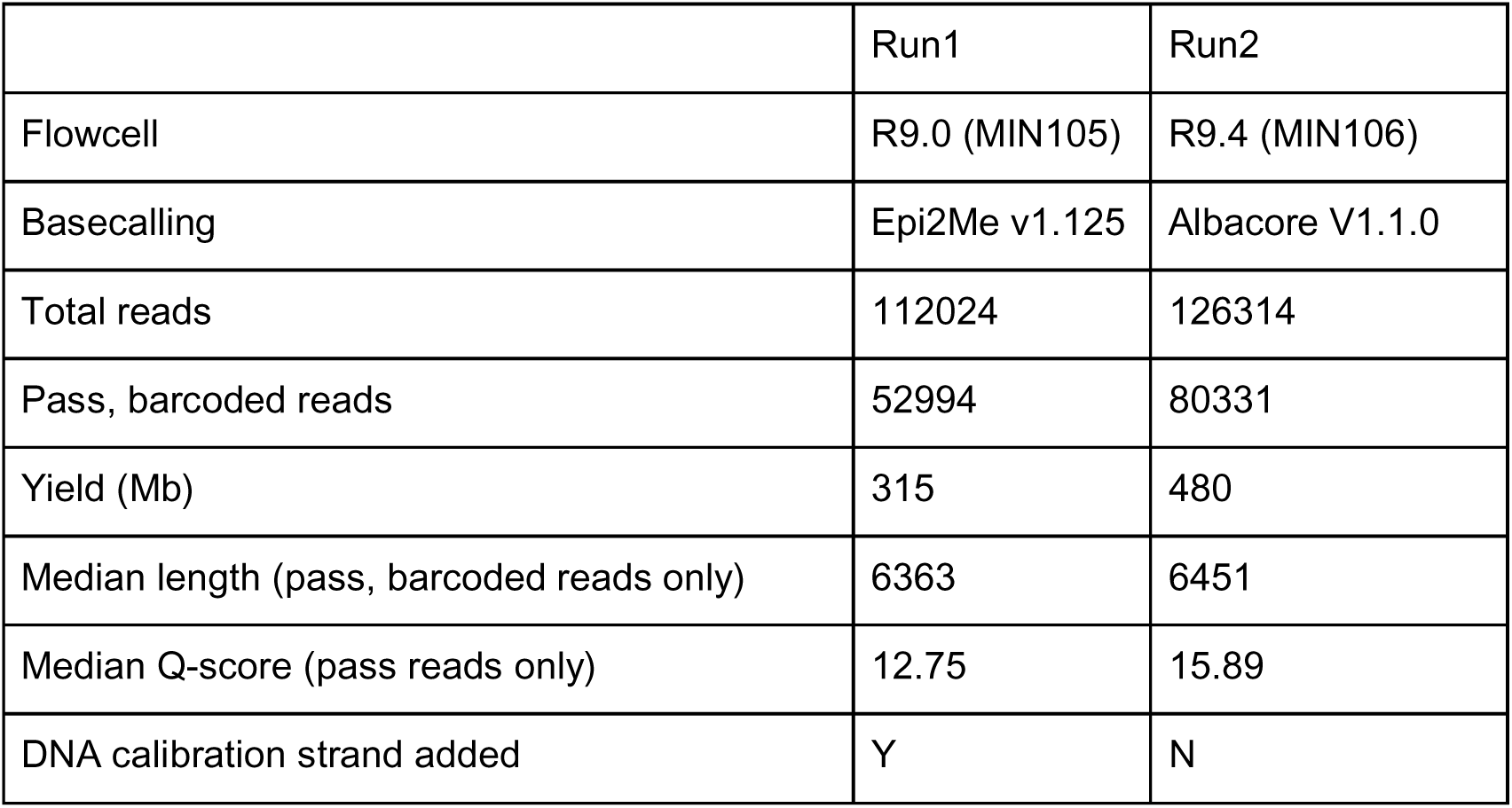
Sequencing metrics for each nanopore sequencing run

**Supplementary Figure 1:**
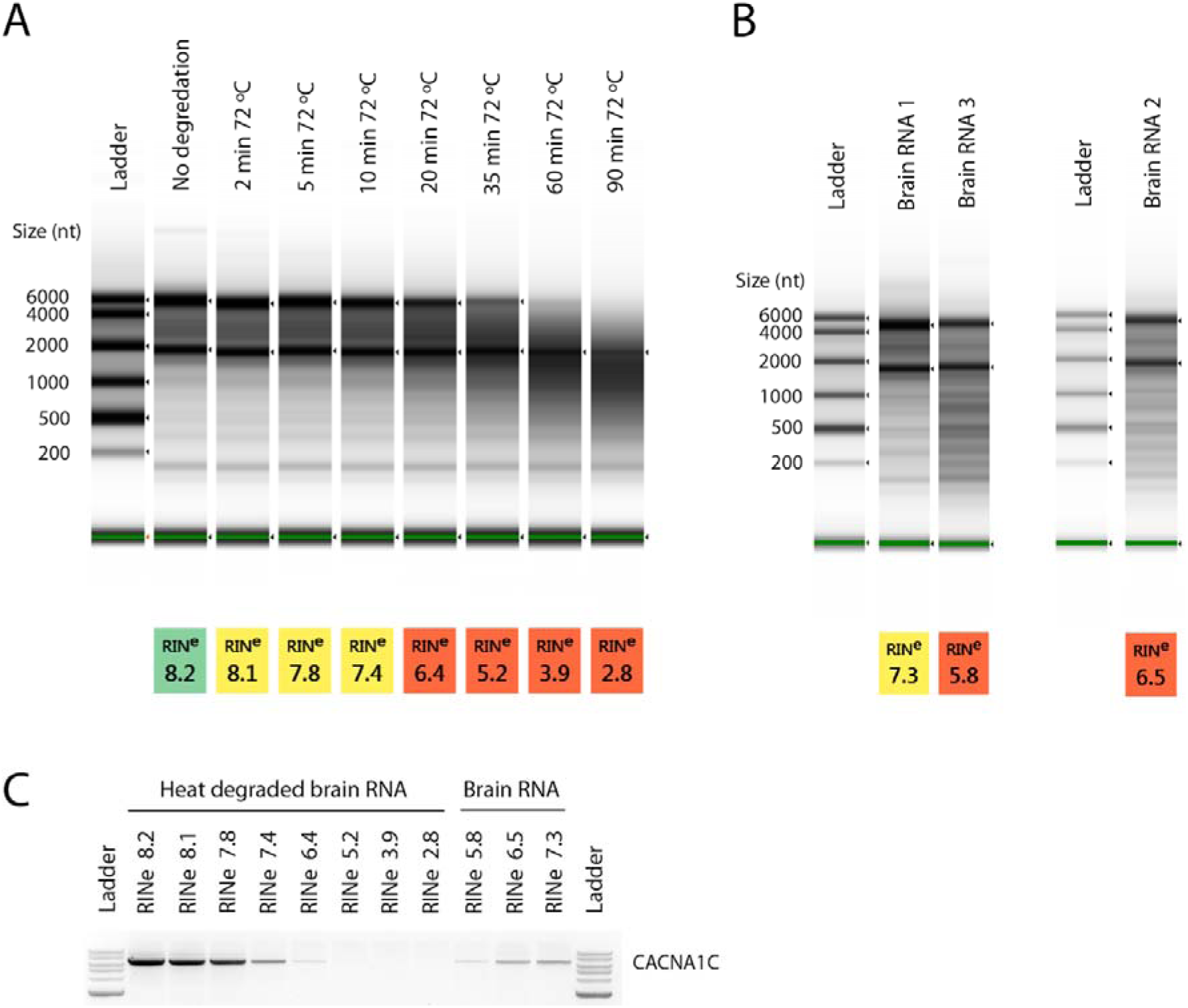
High-quality RNA is required for amplification of *CACNA1C* CDS. The quality of RNA from post-mortem or tissues samples can be highly variable with many samples having undergone significant degradation. Conversely, RT-PCR and sequencing of long and/or full-length cDNAs requires the sample RNA to be of sufficient integrity to contain undegraded transcripts. To investigate how RNA quality impacts the feasibility of amplifying and sequencing full-length genes, and to establish a minimum recommended quality value, we artificially degraded brain RNA to create a series of RINe values and investigated the effect of RNA quality on full-length CACNA1C CDS amplification. RNA quality varied from RINe 8.2 (undegraded RNA) to 2.8 (90 min at 72°C) (Figure S1A. *CACNA1C* amplified strongly in samples with a RINe of ∼8, with lower quality samples showing decreasing amounts of *CACNA1C* and no product from samples with a RINe of ∼<5 (Figure S1C). As RNA degradation from heat treatment may not accurately reflect standard sample degradation, we attempted *CACNA1C* amplification in three untreated striatal samples with RINe of 7.3, 6.5 and 5.8 with similar results (Figure S1B,C). These results suggest a minimum RINe value of 6 for generating long amplicons from post-mortem brain samples and a recommended RINe of >7 for robust amplification. (**A**) Tapestation profile and RINe quality values of brain RNA after heat treatment at 72°C. Untreated RNA had a RINe of 8.2. (**B**) Tapestation profiles and RINe quality values of untreated striatal brain RNA samples. Only lanes of interest from Tapestation profiles are shown. (**C**) Agarose gel showing amplification of the ∼6.5kb *CACNA1C* CDS from RNA of varying quality.

**Supplementary Figure 2:**
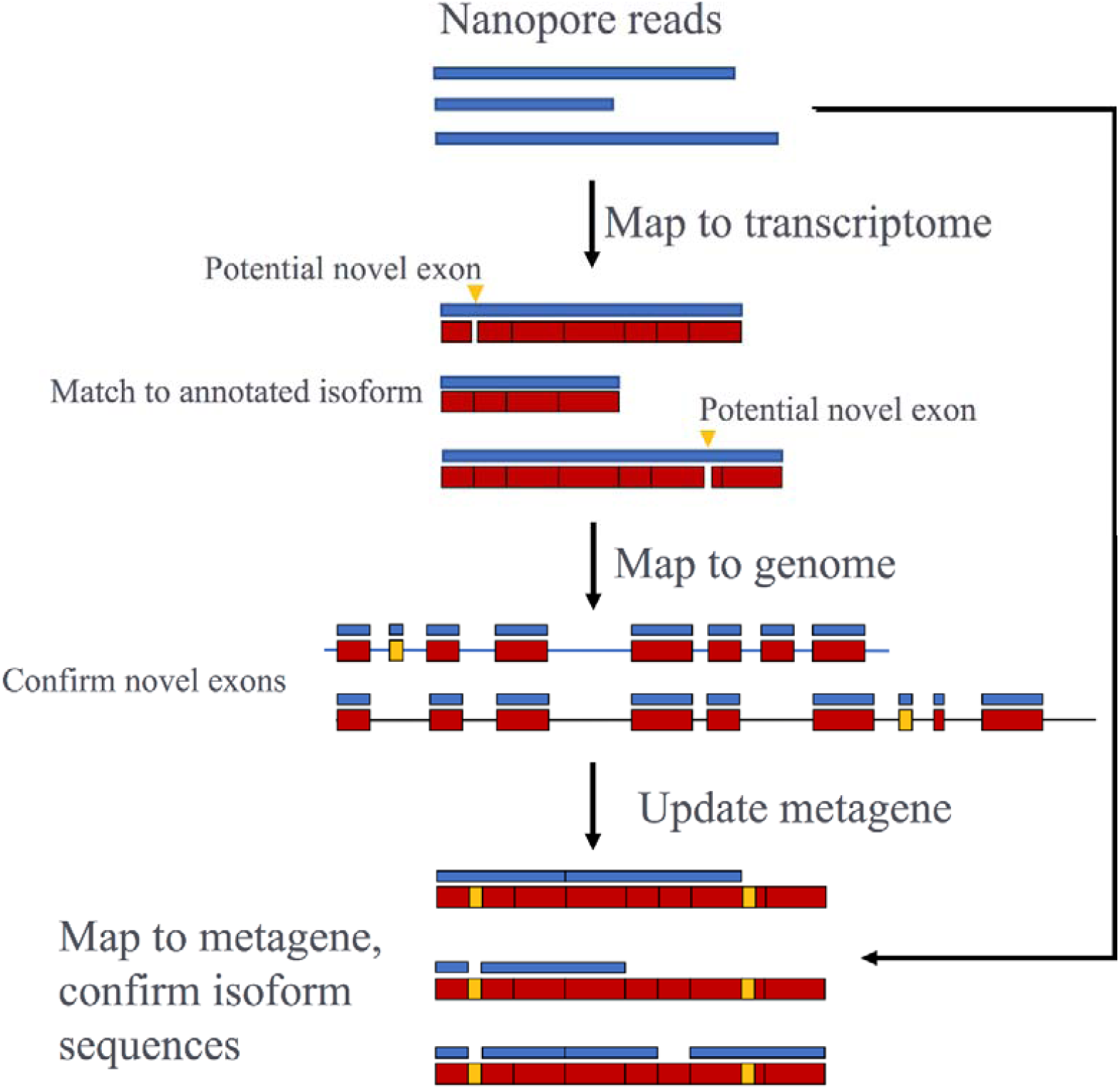
**Mapping pipeline for the annotation of novel exons and novel transcripts**

**Supplementary Figure 3:**
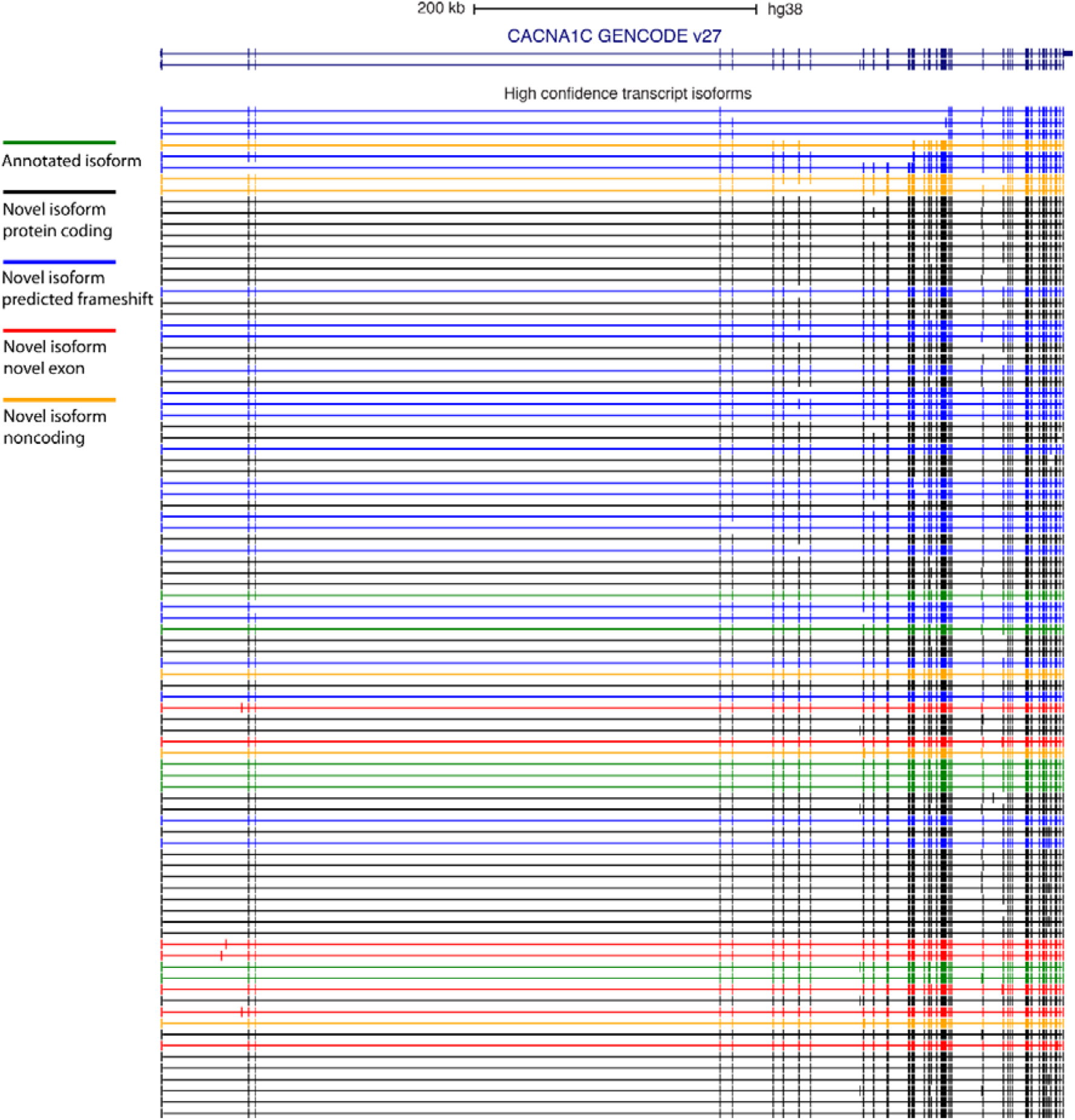
**High confidence *CACNA1C* isoforms identified by Nanopore sequencing** USCS genome browser image of all high confidence CDS transcripts from the exon-level analysis. Representative annotated transcripts from GENCODE v27 shown. Colours of nanopore identified transcripts represent different transcript classes based on their coding potential. Two novel exon transcripts (red) are further annotated as noncoding, the remainder as coding.

**Supplementary Figure 4.**
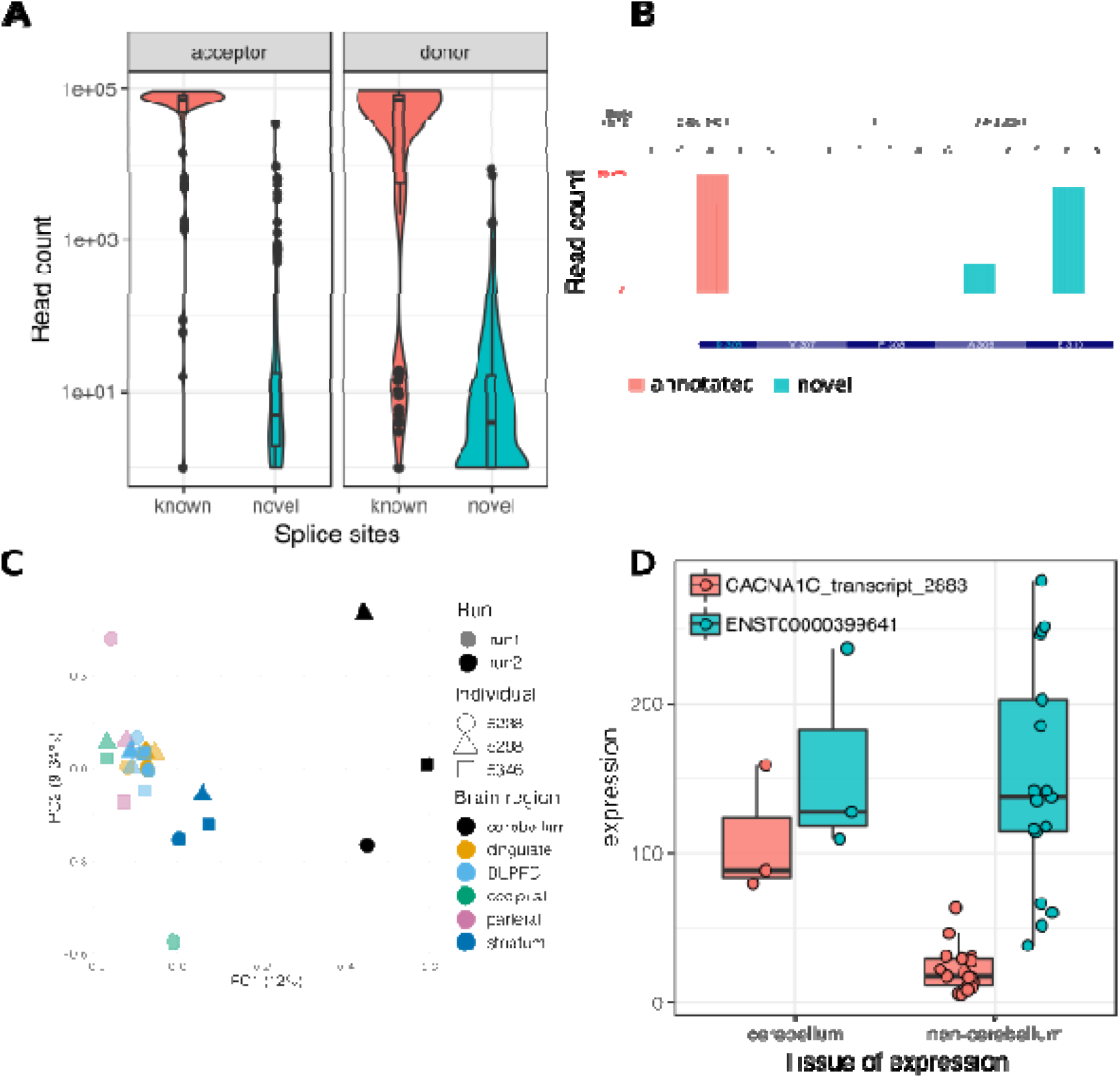
(A) Comparison of annotated and novel splice sites read coverage. (B) Identification of highly supported novel splice sites. (C) Principal Component Analysis based on normalised transcript expression after integration of updated splice site annotation. (D) Isoform switching of *ENST00000399641* and *CACNA1C n2883* (known as n2199 in the exon-level analysis) in cerebellum.

**Supplementary Figure 5:**
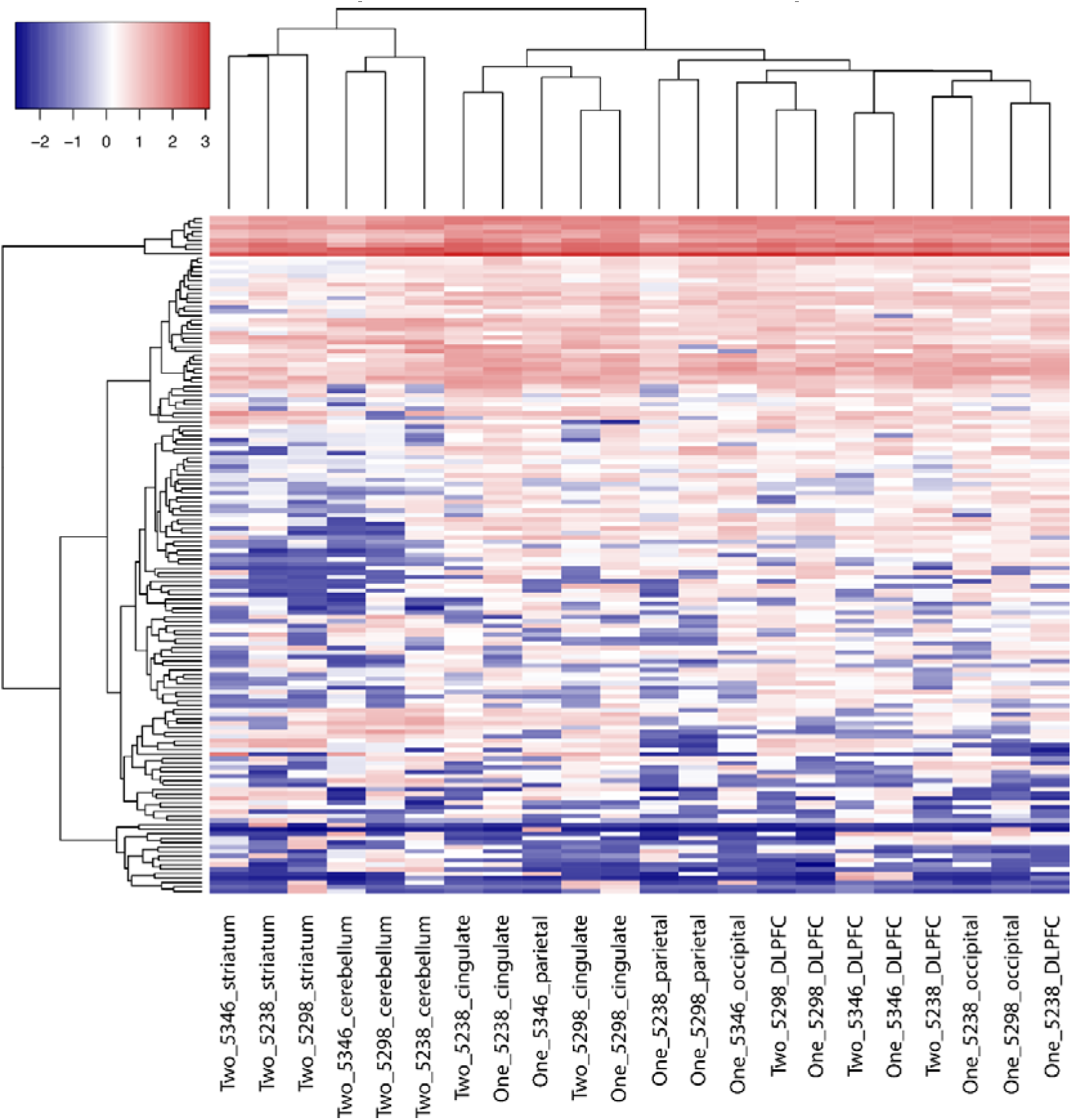
**Clustering of *CACNA1C* isoform expression between individuals and tissues with the permissive set of filtered transcripts** Transcript expression levels across tissues and individuals. “One” and “Two” denote sequencing runs. Due to the large difference in sequencing depth between samples, all libraries were downsampled to the smallest sequencing depth. Inclusion in “permissive” transcript set required isoform identification in at least 2 libraries with a minimum of 24 reads in total. Permissive set of isoforms demonstrates improved separation between brain regions.

**Supplementary Figure 6:**
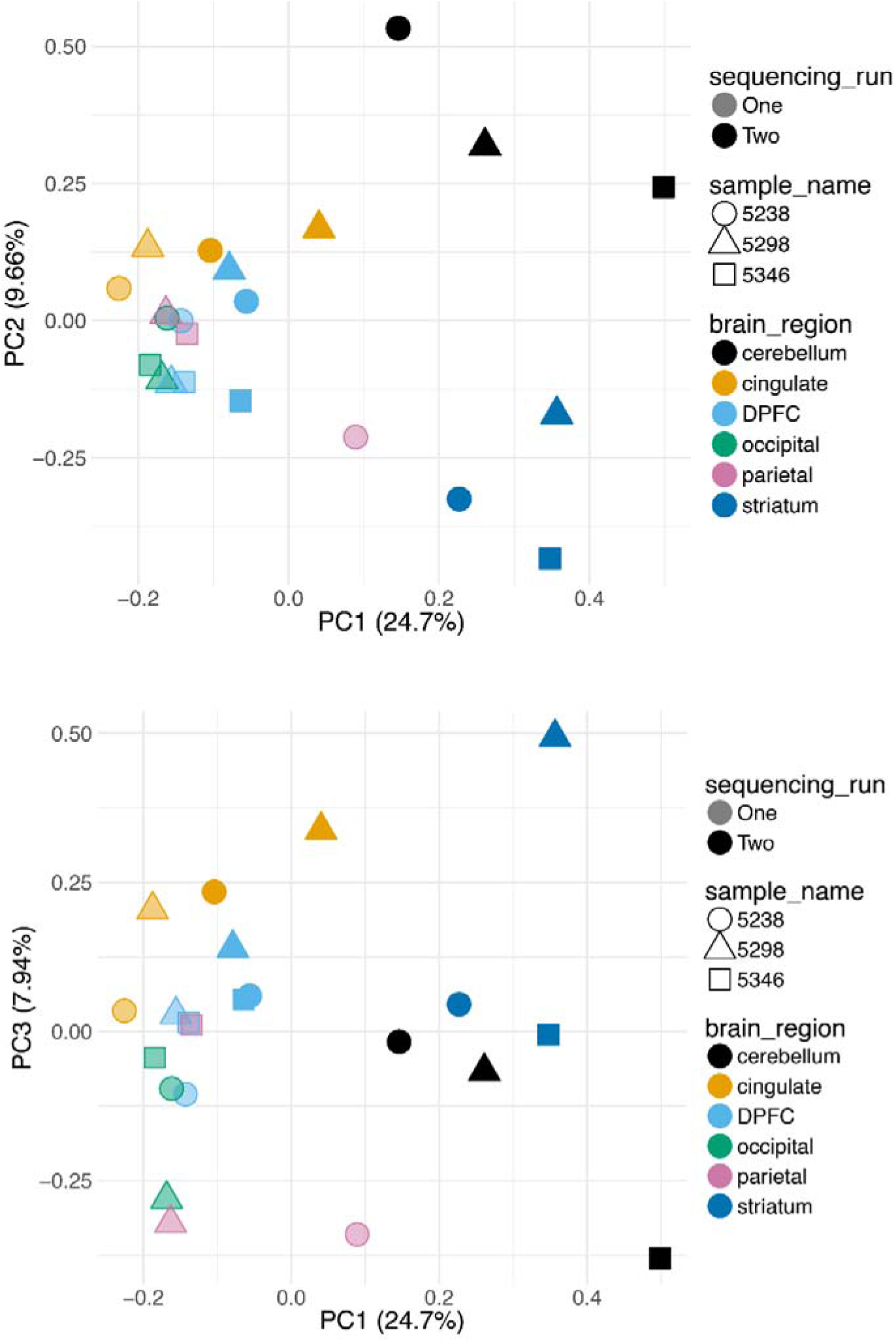
**Principal Component Analysis of *CACNA1C* isoform expression between individuals and tissues with the permissive set of filtered transcripts** Principal Component Analysis based on normalised transcript expression. Due to the large difference in sequencing depth between samples, all libraries were downsampled to the smallest sequencing depth. Inclusion in “permissive” isoform set required transcript identification in at least 2 libraries with a minimum of 24 reads in total. Permissive set of transcripts demonstrates improved separation between brain regions.

**Supplementary Figure 7.**
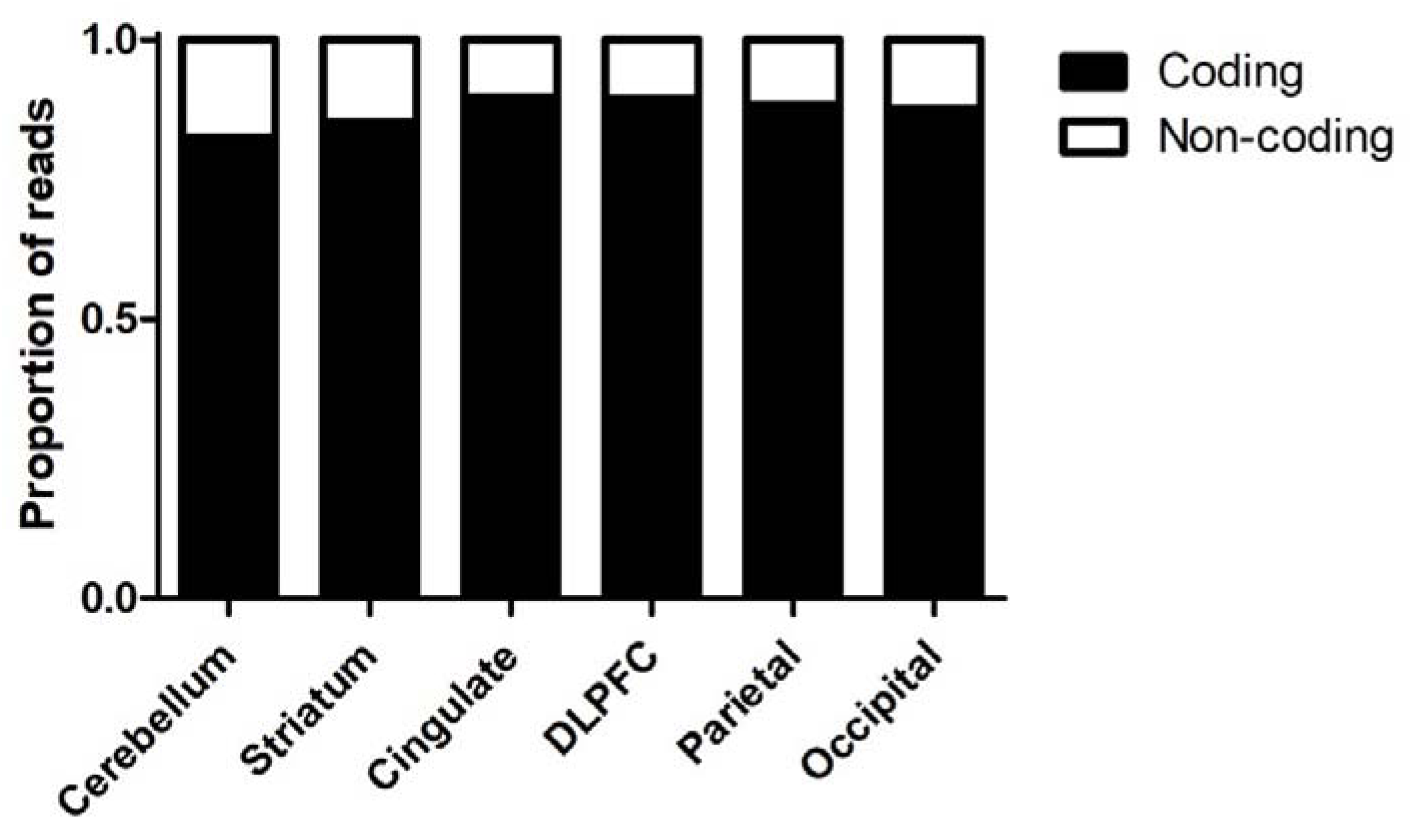
The majority of reads map to *CACNA1C* transcripts that predicted to encode a functional protein. The figure shows the proportion of *CACNA1C* reads that are predicted to encode a functional protein (black bars), compared to those which introduce a frame-shift or which include sequence variants predicted to disrupt transmembrane regions (‘non-coding; white bars) in the different brain regions studied.

**Supplementary Figure 8:**
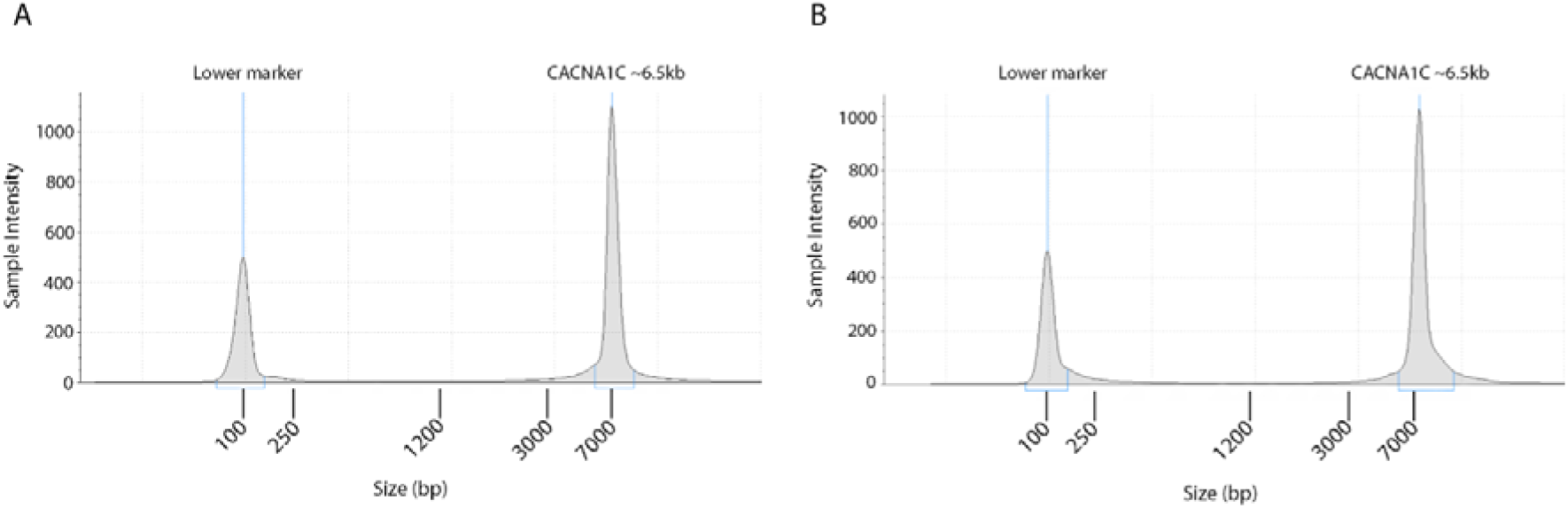
**Full-length, highly pure pooled CACNA1C amplicons.** A) Run1 B) Run2. Slight apparent shift of amplicon to a larger size is due to slight inaccuracy inherent in gDNA screentape where there is only a lower marker for sizing.

## Supplementary Data Files

*Supplementary Data 1:*

Details of novel exons in .bed format.

*Supplementary Data 2:*

Identity of low confidence transcripts from exon-level analysis in .bed format.

*Supplementary Data 3:*

Abundance of low confidence transcripts from exon-level analysis in different samples.

*Supplementary Data 4:*

Identity of high confidence transcripts from exon-level analysis in .bed format.

*Supplementary data 5:*

Abundance of high confidence transcripts from exon-level analysis in different samples.

*Supplementary Data 6:*

Identity of splice-site analysis transcripts in .bed format.

*Supplementary Data 7:*

Abundance of splice-site analysis transcripts from exon-level analysis in different samples.

